# Real-time, MinION-based, amplicon sequencing for lineage typing of infectious bronchitis virus from upper respiratory samples

**DOI:** 10.1101/634600

**Authors:** Salman L. Butt, Eric C. Erwood, Jian Zhang, Holly S. Sellers, Kelsey Young, Kevin K. Lahmers, James B. Stanton

## Abstract

Infectious bronchitis (IB) causes significant economic losses in the global poultry industry. Control of infectious bronchitis is hindered by the genetic diversity of the causative agent, infectious bronchitis virus (IBV), which has led to the emergence of several serotypes that lack complete serologic cross-protection. While serotyping by definition requires immunologic characterization, genotyping is an efficient means to identify IBVs detected in samples. Sanger sequencing of the S1 subunit of the spike gene is currently used to genotype IBV; however, the universal S1 PCR was created to work from cultured IBV and it is inefficient at detecting mixed isolates. This paper describes a MinION-based AmpSeq method that genetically typed IBV from clinical samples, including samples with multiple isolates. Total RNA was extracted from fifteen tracheal scrapings and choanal cleft swab samples, randomly reverse transcribed, and PCR amplified using modified S1-targeted primers. Amplicons were barcoded to allow for pooling of samples, processed per manufacturer’s instructions into a 1D MinION sequencing library, and sequenced on the MinION. The AmpSeq method detected IBV in 13 of 14 IBV-positive samples. AmpSeq accurately detected and genotyped both IBV lineages in three of five samples containing two IBV lineages. Additionally, one sample contained three IBV lineages, and AmpSeq accurately detected two of the three. Strain identification, including detection of different strains from the same lineage, was also possible with this AmpSeq method. The results demonstrate the feasibility of using MinION-based AmpSeq for rapid and accurate identification and lineage typing of IBV from oral swab samples.

## Introduction

Infectious bronchitis (IB), which is caused by infectious bronchitis virus (IBV), is one of the most important diseases of poultry, causing severe economic losses worldwide.^8^ Clinical signs of disease are diverse and include respiratory distress, severe ocular discharge, poor body weight gain, decreased egg production, flushing (renal disease), and occasionally mortality in chickens.^7^ IB is often complicated by secondary bacterial (e.g., *E. coli, Mycoplasma* sp.) and viral infections (e.g., avian influenza virus, Newcastle disease virus, and infectious laryngotracheitis virus).^43^ Lack of cross protection among IBV serotypes is a challenge to controlling IB^15^; therefore, control of IB relies heavily on serotype-specific live attenuated vaccines.^8^ Collectively, the presence of multiple IBVs in a single sample, emergence of variant IBVs, and the high genetic diversity of IBV can complicate the diagnosis of IB and illustrate the need for enhanced diagnostics.^15^

Infectious bronchitis virus is an enveloped, pleomorphic gammacoronavirus with an unsegmented, single-stranded, positive-sense, 26-27.8 Kb, RNA genome that encodes the non-structural polyprotein,1a and 1b, and several structural proteins: spike (S), envelop (E), membrane (M) and nucleocapsid (N).^38,41^ In addition, two accessory genes, expressing 3a, 3b and 5a and 5b, respectively, have also been described.^6,14,38^ The S protein is highly glycosylated and post-translational cleavage leads to two subunits: S1 and S2.^10,48^ Besides acting as the viral attachment protein, the S1 protein is a major target of neutralizing antibodies.^7^ As with many attachment proteins that are targets of virus-neutralizing antibodies, the S1 subunit is highly diverse with almost 50% of the amino acids differing among IBV serotypes.^2,21,38^ Such variation leads to important biological differences between IBV serotypes and the emergence of novel variants. More than 60 serotypes of IBV have been reported, but the most common serotypes of IBV in North America are Arkansas, Connecticut, Delaware072, GA08, GA98 and Massachusetts.^16^ This genetic diversity leads to emergence of new serotypes and a lack of complete cross-serotype protection by vaccines.^29^ The correlation of IBV genotypes and serotypes of IBV has been reported ^45^; therefore, accurate genotypic identification of IBV is an important step to diagnose IBV in clinical respiratory cases, ensure selection of proper IBV vaccines for use in vaccination programs, and to better understand the epidemiology of this global virus.

One of the recent comprehensive classification schemes for IBV uses S1 gene sequence-based phylogeny of IBV and identified six genotypes (I-VI), 32 sub-genotypic lineages, and a number of inter-lineage recombinants in global strains of IBV.^44^ Among the six genotypes, genotype I (G-I) is the most diverse group of viruses, with 27 unique lineages.^44^ As such, sequencing of the S1 subunit provides important information regarding the classification of IBV within samples.

The genetic classification of IBV has relied on genotype-specific RT-qPCR assays, serotype-specific S1 RT-qPCR ^32,40^ and/or pan-IBV S1 RT-PCR assays coupled with Sanger sequencing.^8,31,49^ Genotype-specific RT-qPCRs are limited to short fragments, which may miss important changes in the S1 (~1.6 kb)^44^ outside the short target. Additionally, since the target is short, the primers lie within the variable regions of the S1 and may require a different assay for each genotype. Lastly, the pan-IBV S1 primers^1,17,22,23,25,33^, while only requiring one reaction, have relatively low sensitivity and are typically used only on egg-cultured virus, which adds an additional, time consuming step and many diagnostic labs do not have SPF embryos readily available. Lastly, cloning of PCR products has been used to detect multiple IBV sub-populations in the sample^30^; however, this is inefficient when potentially dealing with multiple (three or four) IBV types. As such, the development of a pan-IBV sequencing method to rapidly determine the genotype(s) in samples would aid in lineage typing and tracking of IBV genotypes circulating in poultry flocks.

Recently, third generation sequencing technology has been used for detection of viral nucleotides and sequencing ultra-long DNA molecules.^34^ The MinION Nanopore sequencer, a new DNA sequencing technology that allows for rapid, in-house, real-time technique detection and differentiation of IBV lineages, may be cost effective and useful in the field.^19^ Amplicon-based sequencing has also been used to amplify specific regions of Newcastle disease^3,13^, infectious laryngotracheitis virus^39^ Zika^34^, Ebola^35^ and avian influenza viruses^46^, by simple RT-PCR and then sequencing on the MinION device. Real-time data analysis, the lack of significant start-up cost investment or maintenance expenses, simultaneous and sequential multiplexing unique to MinION, and the ability to sequence long DNA molecules so that primers are in conserved regions while the product contains the variable region are the features that make this technology highly feasible to be used for disease diagnostics.

Accurate identification of IBV genotypes from samples, including detection of multiple isolates from a single sample, is crucial to respiratory disease diagnosis, selection of appropriate IBV vaccines, and epidemiologic studies. Therefore, the aim of the current study is to create a single, amplicon-based protocol to sequence IBV-S1 gene and develop a sequence analysis workflow to identify IBV isolates from clinical swab samples. This method provides a useful assay for IBV and a model for the development of future amplicon sequencing (AmpSeq) based assays.

## Material and methods

### Samples

Clinical swab samples (n = 15) were obtained from samples submitted to the Poultry Diagnostic & Research Center, University of Georgia. Detailed information about the samples is provided in Table 1.

**Table 1.** Background information of clinical samples collected from broiler chickens.

| Assigned IDs | Sample | Run | Study | Pan-IBV RT-qPCR (Cq)* | GA08 specific RT-qPCR (Cq) |
| --- | --- | --- | --- | --- | --- |
| 1 | B2-42 | 1 | Vaccine study | 30.90 | 38.02 |
| 2 | B2-33 | 1 | Vaccine study | 24.27 | 29.45 |
| 3 | B2-25 | 1 | Vaccine study | 25.46 | Neg |
| 4 | B2-24 | 1 | Vaccine study | 27.04 | Neg |
| 5 | B2-21 | 1 | Vaccine study | Neg | Neg |
| 6 | B1-31 | 1 | Vaccine study | NT | 27.00 |
| 7 | B1-10 | 1 | Vaccine study | NT | 24.28 |
| 8 | B1-29 | 1 | Vaccine study | NT | Neg |
| 9 | B1-22 | 1 | Vaccine study | NT | Neg |
| 10 | B1-19 | 1 | Vaccine study | NT | Neg |
| 11 | 123539 | 2 | Respiratory disease | 18.83 | NT |
| 12 | 124038 | 2 | Respiratory disease | 19.09 | NT |
| 13 | 123835 | 2 | Respiratory disease | 17.52 | NT |
| 14 | 124686 | 2 | Respiratory disease | 18.09 | NT |
| 15 | 124778 | 2 | Respiratory disease | 19.21 | NT |

NT; not tested
Neg; negative
Cq; quantitation cycle

### IBV RT-qPCR assay

Total RNA was extracted from each of the swab samples using QIAamp Viral RNA Mini kit (Qiagen, Hilden, Germany) as per manufacturer’s instructions, aliquoted, and stored at −80 °C until further use. These samples were previously tested for IBV and GA08 serotype of IBV. Briefly, a pan IBV RT-qPCR assay^4^ was used to detect IBV in general and then IBV strain specific RT-qPCR assay^36^ was used to detect GA08 in the samples using the AgPath ID, One step RT-PCR kit (Thermo Fisher Scientific, Waltham, MA, USA) on the Applied Biosystems 7500 FAST following the previously described protocols.

### MinION cDNA synthesis

For MinION cDNA synthesis, RNA was extracted from 500 μ1 of each clinical sample using Trizol-LS (Thermo Fisher Scientific, Waltham, MA, USA), per manufacturer’s directions. A reaction mixture of 8 μ1 of total RNA, 1 μ1 of random primers and 1 μ1 of dNTPs was incubated at 65 °C for 5 min, chilled on ice for at least 1 min followed by addition of 10 μ1 of cDNA synthesis mix including SuperScript III (Thermo Fisher Scientific, Waltham, MA, USA) according to the manufacture’s instruction. The reaction was incubated at 25 °C for 10 min, then at 50 °C for 50 min for cDNA synthesis. The reaction was terminated at 85 °C for 5 min, and then chilled on ice. To remove residual RNA, the cDNA solution was incubated with RNase H at 37 °C for 20 min.

### MinION amplicon synthesis

A universal S1 primer set,^27^ tailed with the MinION universal adapter sequence of 22 nucleotides (underlined, Table 2) to allow barcoding of amplicons, was used for targeted amplification of the IBV S1 gene for IBV. The PCR reaction mixture (Expand High Fidelity PCR system, Roche Diagnostics, Mannheim, Germany) was composed of 10 μ1 of cDNA, 1 μ1 of 10 μM forward primer, 1 μ1 of 10 μM reverse primer, 2.5 μ1 of 10x Expand high fidelity buffer, 1 μ1 of Expand High Fidelity Enzyme Mix, and 1 μ1 of 10mM dNTPs, to a final volume to 25 μ1 with 9 μ1 of nuclease-free water. The following thermocycling conditions were used for amplicon synthesis: denaturation at 95 °C for 2 min; 30 cycles of 94 °C for 1 min, 45 °C for 2 min, and 72 °C for 2 minutes, and final extension at 72 °C for 10 min. See Table 2 for details about all three primer sets used in this study. PCR products were electrophoresed in 1.5% agarose with SYBR Safe DNA gel stain (Invitrogen) for visual evaluation. Amplified DNA was purified by Agencourt AMPure XP beads (Beckman Coulter, USA) at 1.6:1 (volumetric bead:DNA) and quantified using the dsDNA High Sensitivity Assay Kit on a Qubit^®^ Fluorimeter 3.0.

**Table 2.**
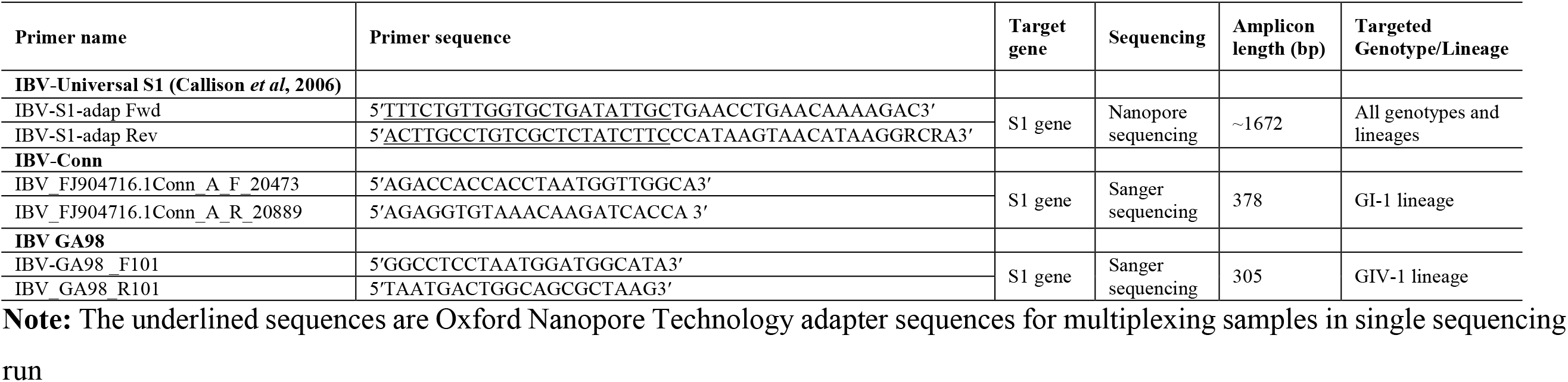
Details of PCR primer sets used to detect IBV in samples.

### Library preparation and MinION Sequencing

The amplicons obtained from the tailed S1 primer set were then used to prepare MinION-compatible DNA libraries. Briefly, each of the amplicons were diluted to 0.5 nM and amplified using barcoding primers (1D PCR Barcoding Amplicon Kit, Oxford Nanopore Technologies, [ONT], UK) and LongAmp Taq 2X Master Mix (New England Biolabs, MA, USA) with the following conditions: 95 °C for 3 min; 15 cycles of 95 °C for 15 seconds, 62 °C for 15 seconds, 65 °C for 80 seconds; and 65 °C for 80 seconds. The barcoded amplicons were bead purified (1:1.4; bead:solution), pooled (Run1; pool of 10 samples, Run2; pool of 5 samples) into a single tube, end prepped with dA tailing Kit (NEBNext^®^ Ultra™ End Repair/dA-Tailing Module, New England Biolabs, MA, USA), bead purified (1:1; bead:solution), and ligated to the sequencing adapters (Ligation Sequencing Kit 1D, SQK-LSK108, ONT, UK), all per Oxford Nanopore Technology directions. Final DNA libraries were bead purified (0.4:1; bead:solution), eluted in 15 μL and sequenced with the MinION Nanopore sequencer. A new FLO-MIN106 R9.4 flow cell (ONT), stored at 4°C prior to use, was allowed to equilibrate to room temperature for 10 minutes and then primed with running buffer as per manufacturer’s instructions. The pooled DNA libraries were prepared by combining 12 μL of the library pool with 2.5 μL nuclease-free water, 35 μL RBF, and 25.5 μL library loading beads. After the MinION Platform QC run, the DNA library was loaded into the MinION flow cell via the SpotON port. The standard 48-h 1D sequencing protocol was initiated using the MinKNOW software v.5.12. Sequencing was allowed to continue for 2 hours until 42,940 reads were obtained for Run1 and 156,000 reads were obtained for Run 2 in 6 hours.

### Building customized BLAST databases for AmpSeq analysis

First, a lineage-typing database, containing 32 IBV S1 gene sequences (one sequence from each of 32 lineages in the 6 genotypes^44^) and the chicken genome (GCF_000002315.4_Gallus_gallus-5.0), was constructed (referred to as “IBV-lineage-typing database” in this study). A second database was constructed with all the Avian Coronaviruses S1 sequences (n = 7328) available in NCBI (as of: 09-08-2017) and the chicken genome (referred to as “All-IBV database” in this study). Prior to database construction, all sequences were dustmasked (NCBI C++ ToolKit), and each isolate was assigned a unique taxonomy ID as a species hierarchically under the genus *Gammacoronavirus* to allow easy sorting. The local BLAST databases were compiled using default settings. The IBV-lineage-typing database was used to assign IBV lineages to each read, as appropriate. The All-IBV database was used to assign IBV isolates taxonomic ID to each of IBV reads from within IBV lineage read clusters. The IBV-lineage-typing database, which contains only one sequence per lineage, is required as centrifuge divides the score for any given read by the number of hits that have an equal score. Since some lineages are overrepresented in GenBank (e.g., Mass strains), this results in the scores of those reads (e.g., Mass reads) being divided by a large number, effectively reducing the score for any single alignment. However, because there is only one read per lineage in the IBV-lineage-typing database, it has insufficient diversity to determine if there is more than one strain of the same lineage present in a sample. Thus, the All-IBV database provides the diversity required for that analysis.

### AmpSeq IBV lineage and isolate identification

A schematic diagram of the workflow of MinION data analysis is presented in Fig. 1. For Nanopore sequencing data, pre-processing steps were performed to prepare data for downstream analysis. Briefly, Nanopore reads (FAST5) were basecalled using Albacore v 2.02 (https://github.com/Albacore/albacore) with following parameters (read_fast5_basecaller.py -i /Input_reads.fast5 --recursive -t 4 -s /Output_files --flowcell FLO-MIN107 --kit SQK-LSK108 – o fastq). Porechop (https://github.com/rrwick/Porechop) was used for adapter trimming, (default setting), barcode-based demultiplexing (default settings), and to trim an additional 21 nucleotides, representing the S1 primer sequences (porechop -i Input_file.fastq --extra_end_trim 21 –b ./output_demultiplexed files.fastq). After barcode and adapter removal, reads were analyzed with a script-based, 2-step data analysis protocol, which includes Centrifuge-kreport as taxonomic read classifier^24^ using the sequences in the above mentioned BLAST databases. Briefly, basecalled reads (FASTQ) from individual barcoded samples were used as an input. First, the basecalled reads were aligned to the IBV-lineage-typing database using BLASTn and reads were clustered based on the read sequence alignment to the respective prototype sequence of IBV lineage. These read clusters were used to interpret presence of IBV genotypes and lineages in the samples. For isolate level identification of IBV, the lineage-based read clusters were individually aligned to the All-IBV database, which produced sub-clusters of reads. Each of these read sub-clusters potentially represents a different isolate and was further used for interpretation. Knowing that MinION sequencing has a high sequencing error rate in individual reads, further steps were added in the data analysis algorithm to obtain a more accurate consensus sequence. Therefore, these read sub-clusters were mapped, using Geneious mapper in Geneious software v11.1.3. (Biomatters, Ltd, Auckland, New Zealand), to the IBV-lineage-typing database (a reference FASTA file) to obtain consensus sequences from each sub-cluster. A minimum threshold for number of reads per consensus sequence was not set as in two samples (sample #4 in Run1 and sample #12 in Run 2), only 1 and 5 IBV reads were obtained, respectively; therefore, consensus sequences were built from all the available IBV reads per subcluster. The consensus sequences were compared to GenBank using BLASTn. To select the “top hit” from BLASTn output, sequence search results were ordered by “sequence identity” and then sequence alignments were evaluated for “minimum mismatches” and “coverage” of query or subject sequence. All isolates with the highest query or subject coverage and the fewest mismatches were used as “top hit” in the final results.

**Figure 1.**
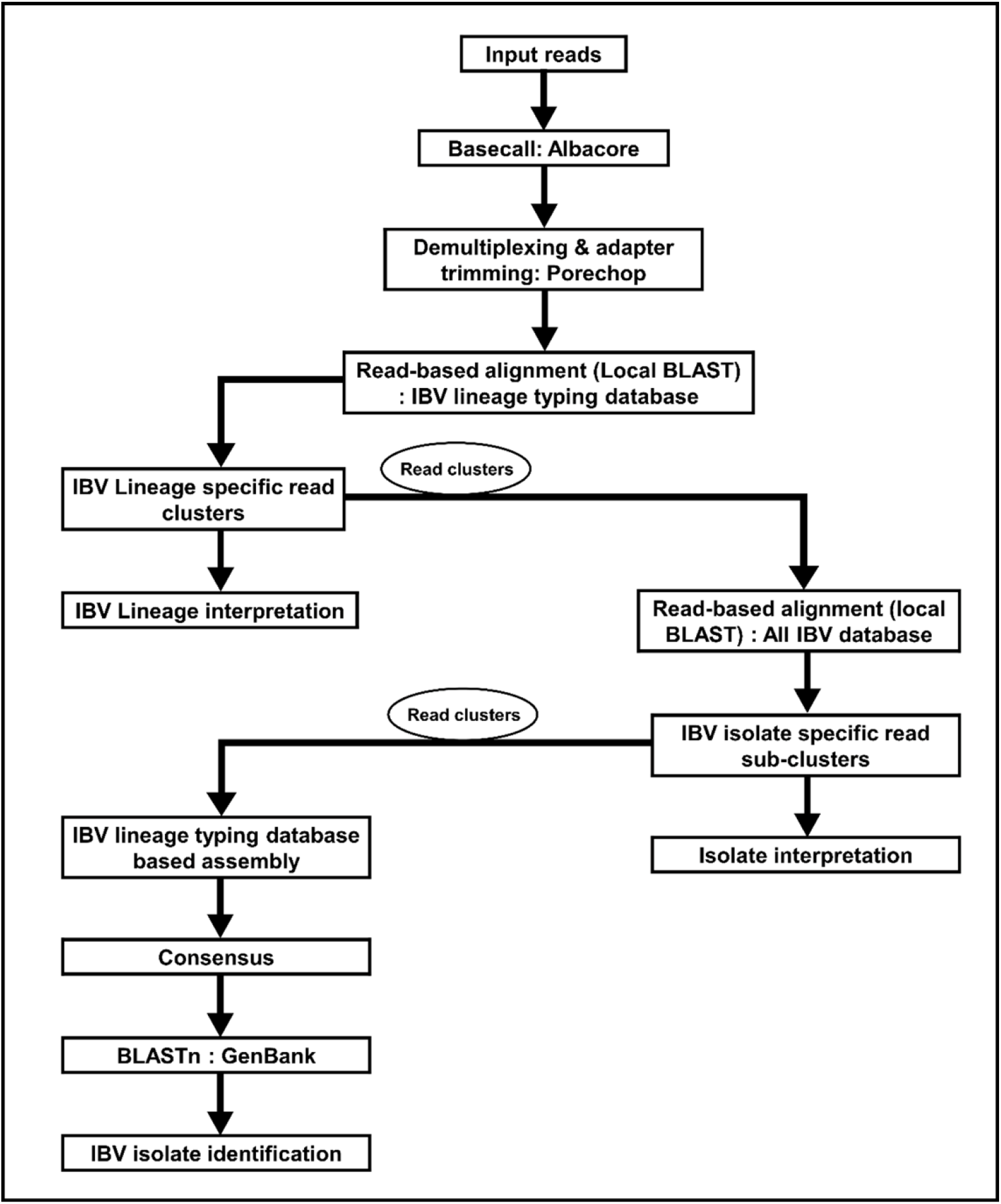
Schematic diagram of workflow for MinION sequence data analysis.

### Sanger sequencing amplicon synthesis

For Sanger sequencing amplicons synthesis, cDNA from MinION library preparation was amplified using the following primer sets (Table 2). Firstly, a primer set based on Connecticut sequence NCBI FJ904716.1 was used to amplify Genotype I-lineage 1 viruses (e.g., Connecticut and Massachusetts serotypes). Secondly, a primer set based on GA98 sequence NCBI AF274439.1 was used to amplify Genotype IV-lineage 1 viruses (e.g., Georgia 1998 and Delaware 072 serotypes). Primer sets were designed using NCBI Primer-BLAST.^51^ The PCR reaction mixture is composed of 10 μ1 of cDNA, 0.5 μ1 of 10 μM forward primer, 0.5 μ1 of 10 μM reverse primer, 2.5 μ1 of 10x buffer, 0.5 μ1 of enzyme mix, and 0.5 μ1 of 10mM dNTPs, then made the final volume to 25 μ1 with nuclease-free water. The following thermocycling conditions were used for amplicon synthesis: denaturation at 95 °C for 1 min, followed by 35 cycles of 94 °C for 30 s, 55 °C (ConnA primer set) or 50 °C (GA98 primer set) for 30 s, and 72 °C for 45 sec, and a final extension of 72 °C for 5 min. PCR products were visually inspected after electrophoresis in 1.5% agarose gel and the correctly sized bands were cut out for cDNA purification using QIAGEN PCR Purification Kit and quantified using the dsDNA High Sensitivity Assay Kit on a Qubit^®^ Fluorimeter 3.0. Briefly, purified amplicons were inserted into plasmids and ligation reactions (10 μ1) were set up as per manufacturer’s instructions (Promega pGEM-T Easy vector system). After ligation, 3 μ1 of ligation mixture was transformed to JM109 competent cells by heat shock. Individual bacterial colonies were checked with PCR and the positive bacterial colonies were plated on LB-agar/ampicillin plates at 37°C for 16 hr. The plasmids were extracted from these positive bacterial colonies (QIAprep Spin Miniprep Kit, Qiagen, Hilden, Germany) and submitted for bidirectional, commercial (GENEWIZ, NJ, USA) Sanger sequencing.

### Sanger sequencing analysis

For Sanger sequencing data, the chromatogram from each of the samples was manually checked and primer sequences were trimmed in MEGA 6.0.^42^ Forward and reverse sequences from multiple clones were aligned using MEGA 6.0 software and consensus sequences were compared to GenBank using Basic Local Alignment Tool (BLAST) (https://blast.ncbi.nlm.nih.gov/) (as of: 12-03-2018) and the top hit from BLASTn output was selected, as described above for MinION sequencing data, to identify the IBV isolate in samples.

### Sanger and MinION sequence pairwise identity

A pairwise nucleotide identity comparison on the partial S1 sequences of IBVs from each of the samples sequenced with Sanger sequencing and MinION sequencing was performed. Briefly, the final AmpSeq consensus sequences from identical IBV types in each of the sample were aligned using ClustalW in MEGA6^42^ and this alignment was used for pairwise nucleotide identity using MEGA6.^42^

## Results

### RT-qPCR assays

Five samples from Run 1 were tested with a pan-IBV RT-qPCR assay^4^, and four of those were positive for IBV. Samples with pathogenic IBV strains (Run 2, n = 5) were tested with the pan-IBV RT-qPCR and all five samples were IBV positive (Table 1). Next, a GA08 serotype specific primer set (in house validated set of primers used for diagnostics) was used on all samples from Run 1 (including five samples that were previously tested by the pan-IBV PCR) and four of 10 samples were positive. Sample #3 and #4, which were positive with the pan-IBV RT-qPCR assay, were negative with GA08 specific RT-qPCR assay indicating that these samples contained IBV serotypes other than GA08 and required further testing. Sample #5 tested negative with both assays (Table 1).

### Sanger sequencing

A total of 10 clinical swab samples (Run 1) were processed for Sanger sequencing to confirm the presence of IBV lineages. Based on the MinION results, primers targeting GIV-L1 and GI-L1 (GA98 and Conn and primer sets, respectively) were created. The PCR products were cloned and multiple (6–24) clones from each sample were submitted for Sanger sequencing and results are summarized in Table 3. Using GIV-L1 serotype primers showed that 9 out of 10 samples were positive with 1 sample (#5) negative for IBV (which is consistent with the pan-IBV RT-qPCR results for this sample). Consensus sequences from multiple clones obtained by using GIV-L1 primers showed the presence of GA98 (GA/A9dvaccinated) in 7 of 9 IBV-positive samples. DE072 was detected in samples #1 and #9 as the lone GIV-1 isolate but was also detected in three samples (#2, and #10) that also contained GA98 (Table 4). By using the GI-L1 primer set, which amplified Conn and Mass type IBVs, 3 of 9 IBV-positive samples were positive (#2 and #4 for Conn; #3 for Mass [PDRC_110177]) (Table 4). Based on the two conventional RT-PCRs and the GA08 RT-qPCR, it was determined that only samples #8 and #9 contained a single IBV isolate, while the other 7 samples were positive for 2–4 types of IBV (Table 3).

**Table 3.** Detection of different lineages of IBV in tracheal swab samples using RT-qPCR assays, RT-PCR coupled with Sanger sequencing and MinION sequencing.

| Sample IDs | Lineage IDs |  | Lineage Totals |  |
| --- | --- | --- | --- | --- |
|  | RT-qPCR or PCR with Sanger | MinION-AmpSeq | All 4 RT-PCR assays | MinION-AmpSeq |
| 1 | *GI-L27 \| GIV-L1 | Neg | 2 | 0 |
| 2 | GI-L27 \| GIV-L1 \| GI-L1 | GI-L27 \| GI-L1 | 3 | 2 |
| 3 | GIV-L1 \| GI-L1 | GI-L1 | 2 | 1 |
| 4 | GIV-L1 \| GI-L1 | GIV-L1 \| GI-L1 | 2 | 2 |
| 5 | Neg | Neg | 0 | 0 |
| 6 | GI-L27 \| GIV-L1 | GI-L27 \| GIV-L1 | 2 | 2 |
| 7 | GI-L27 \| GIV-L1 | GI-L27 \| GIV-L1 | 2 | 2 |
| 8 | GIV-L1 | GIV-L1 | 1 | 1 |
| 9 | GIV-L1 | GIV-L1 | 1 | 1 |
| 10 | GIV-L1 | GIV-L1 | 1 | 1 |

\*G-L; Genotype-Lineage
Neg; Negative

### MinION sequencing and lineage identification (Run 1)

PCR amplicons obtained directly from 10 clinical swab samples with vaccine IBV serotypes were barcoded, pooled together and sequenced on MinION device (Run 1). A total of 38,661 reads were successfully basecalled from the entire sequencing run (total reads = 42,940). After demultiplexing, reads per barcode ranged from 831 to 4114. A total of 14,845 reads were not assigned to any of the used barcodes and 128 reads were discarded due to middle adapters. The Nanopore reads were queried against IBV-lineage-typing database to determine if the samples contained IBV (Figure 1). This AmpSeq protocol detected IBV reads in 8 of 10 samples. The number of IBV reads per sample (IBV positive sample) ranged from 56–944. The MinION-negative samples included sample #5, which was consistently negative with RT-qPCR and RT-PCR assays (and thus interpreted as IBV negative), and sample #1, which had the highest Cq values in the pan-IBV RT-qPCR (Table 1).

**Table 4.**
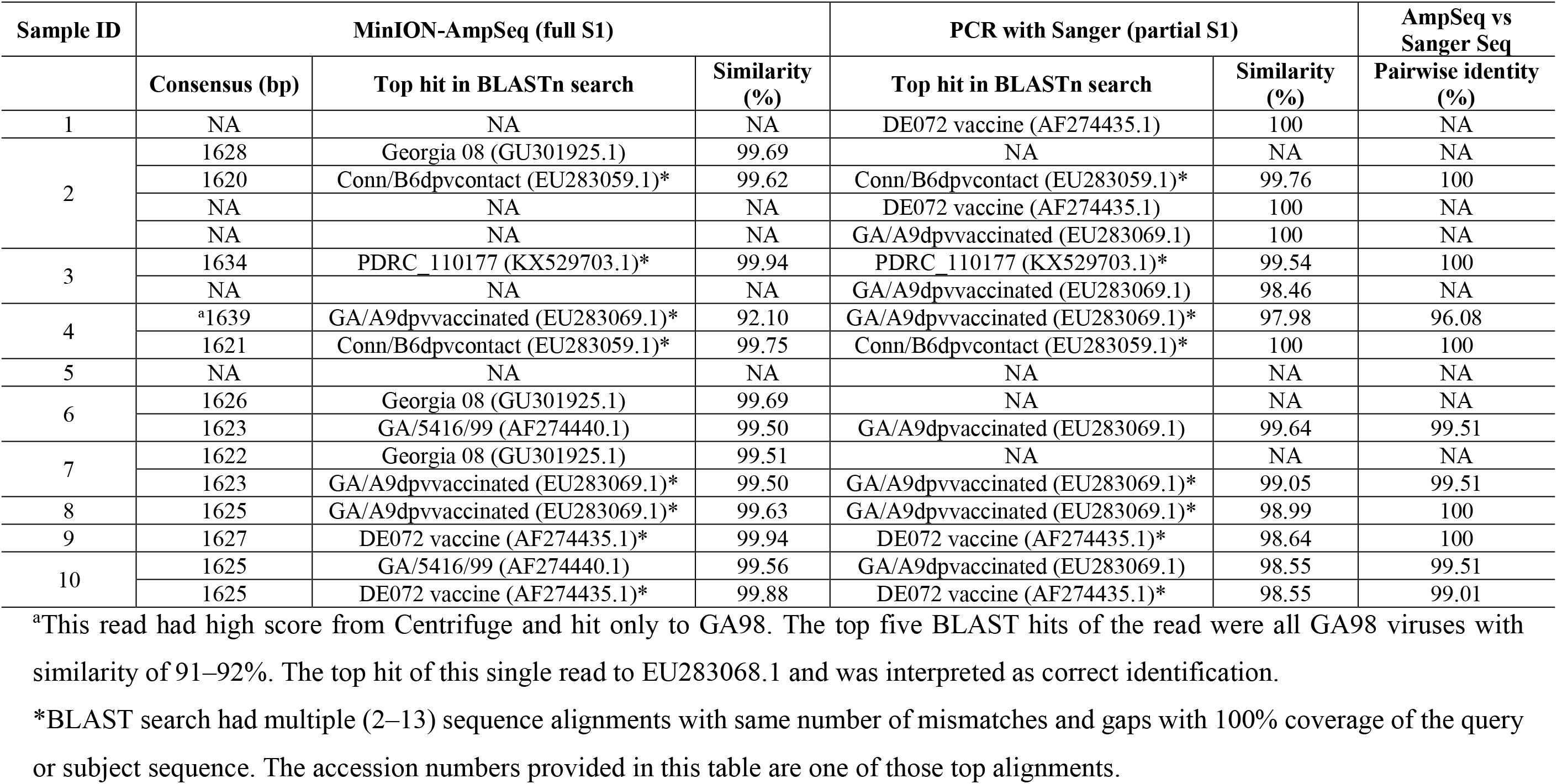
Consensus-based identification of IBV isolates in clinical swab samples

Sequencing data was further analyzed to determine if multiple IBV genotypes or lineages could be detected. Lineage-1 and lineage-27 from Genotype I (GI-L1, GI-L27), and lineage-1 from Genotype IV (GIV-L1) were the detected lineages in the samples. Results from individual RT-qPCR assay for GA08 (GI-L27), and RT-PCR assays for GIV-L1and GI-L1 confirmed the presence of multiple IBV genotypes and lineages in 6 of the 9 IBV-positive samples (#1, #2, #3, #4, #6, #7) and AmpSeq results confirmed multiple genotypes and lineages in 4 (#2, #4, #6, #7) of those 6 samples (AmpSeq failed to detect any IBV in #1, and failed to detect GIV-L1 in #3). The presence of a single lineage was confirmed by PCR-based assays and AmpSeq in three samples (#8, # 9 and #10) (Table 3).

### MinION consensus sequence evaluation for isolate identification

It was observed that the single read cluster composed of less than 5 reads (Run1, sample #4, GA98) yielded a poor-quality consensus sequence, consistent with the known individual read error rate of MinION sequencing. High-quality consensus sequences were obtained from the other read clusters (>5 reads per cluster). Each of the obtained consensus sequences were compared to GenBank databases using BLAST, which revealed >99% sequence identity to respective IBV isolates Table 4). As described in the Sanger Sequencing section, two samples (#8 and #9) contained only one IBV isolate per the non-MinION assays, and the AmpSeq results were consistent with these findings. Non-MinION assays showed that seven samples contained two or more isolates per sample. Six samples (#1, #3, #4, #6, #7, and #10) contained two isolates per sample (Table 4). Of those six samples, AmpSeq detected both isolates in four samples (#4, #6, #7, and #10), one isolate in one sample (#3), and no IBV in one sample (#1; as mentioned above, sample #1 was the sample with the highest Cq value). Of note, sample #10 contained two isolates from the same lineage and AmpSeq was able to identify both isolates within this sample. Finally, the seventh multi-isolate sample (#2) contained four isolates per the non-MinION results, and the AmpSeq detected two of those four isolates. These data show that a co-infection of multiple IBV lineages existed in the above-mentioned samples but a single RT-qPCR, or Sanger sequencing of a single clone may not have detected these co-infecting IBV lineages and required multiple assays to detect all the IBV types. However, the AmpSeq protocol accurately detected multiple IBV lineages in 4 of 7 samples, with partial detection in two of the remaining three samples (Table 4). In samples which had the same IBV type, Blast search of consensus sequences obtained from AmpSeq and Sanger sequencing identified the same (or highly related isolate for sample #6 [GA98] and sample #9) in the NCBI nt database as per parameters described in the methods section.

### MinION sequencing and lineage identification (Run 2)

To evaluate the utility of this protocol on clinical swab samples with pathogenic IBV variants, IBV-S1 gene was amplified directly from five clinical tracheal scrapings and the PCR amplicons were used to create MinION libraries. A total of 146,540 reads were successfully basecalled from the entire sequencing run (total reads = 156,000). After demultiplexing, reads per barcode ranged from 4,285 to 41,131. A total of 24,297 reads were not assigned to any of the used barcodes and 8,109 reads were discarded due to middle adapters in the basecalled reads. Real-time analysis of MinION data, which was obtained within 10 minutes of the sequencing, was sufficient for detection of IBV. However, the sequencing data obtained from entire sequencing run was processed with the same protocol as described above. This AmpSeq protocol detected IBV reads in all of the 5 tested samples using all basecalled reads. The number of IBV reads per sample ranged from 5–4,956. Additionally, the sequencing data analysis showed that the IBV reads belonged to GI-17 lineage and typed the isolate as DMV1639. After MinION sequencing results, these samples were later tested to confirm the presence of IBV variant by MDL_DMV1639 IBV variant specific RT-qPCR assay. All 5 samples were positive for IBV MDL_DMV1639 variant of IBV (Table 5).

**Table 5.**
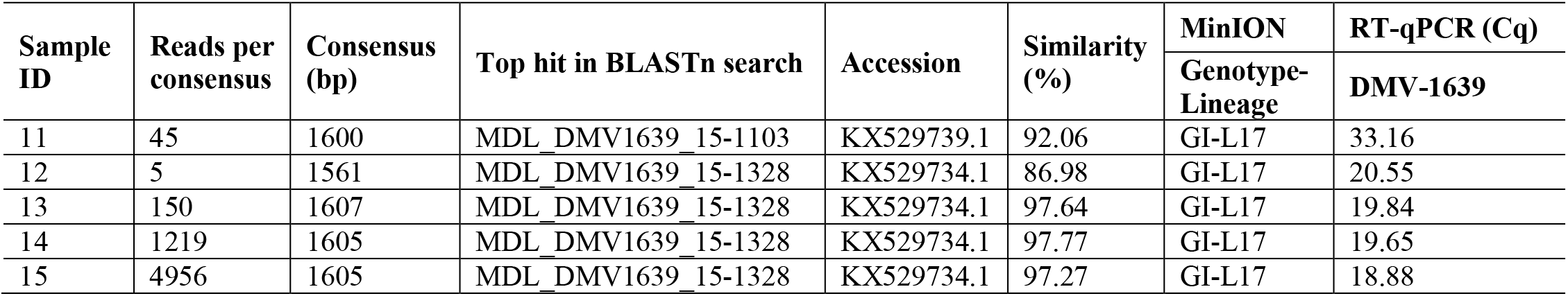
Detection of IBV genotype/lineage using RT-qPCR and MinION AmpSeq in tracheal swab samples (Run 2)

| Sample ID | Reads per consensus | Consensus (bp) | Top hit in BLASTn search | Accession | Similarity (%) | MinION | RT-qPCR (Cq) |
| --- | --- | --- | --- | --- | --- | --- | --- |
|  |  |  |  |  |  | Genotype-Lineage | DMV-1639 |
| 11 | 45 | 1600 | MDL_DMV1639_15-1103 | KX529739.1 | 92.06 | GI-L17 | 33.16 |
| 12 | 5 | 1561 | MDL_DMV1639_15-1328 | KX529734.1 | 86.98 | GI-L17 | 20.55 |
| 13 | 150 | 1607 | MDL_DMV1639_15-1328 | KX529734.1 | 97.64 | GI-L17 | 19.84 |
| 14 | 1219 | 1605 | MDL_DMV1639_15-1328 | KX529734.1 | 97.77 | GI-L17 | 19.65 |
| 15 | 4956 | 1605 | MDL_DMV1639_15-1328 | KX529734.1 | 97.27 | GI-L17 | 18.88 |

### Pairwise sequence identity between Sanger and MinION consensus sequences

In all of the samples where a matching IBV genotype/lineage was identified by both AmpSeq and Sanger sequencing, the AmpSeq data showed high concordance with Sanger sequencing data, In one sample (#4), only a single MinION read of GA98 was detected by AmpSeq and had only 96.08% similar to the respective, shorter sequence from Sanger sequencing, consistent with the reported single-read accuracy of MinION sequencing. All other samples showed 99–100% pairwise identity across the partial S1 fragment generated by Sanger Sequencing. (Table 4).

## Discussion

This study describes a sequencing approach for rapid and accurate identification of IBV genotypes/lineages utilizing the MinION Nanopore sequencer. Additionally, a sequence analysis workflow is described here. The workflow analysis is designed to deal with the multiple, highly similar sequences contained within GenBank that are due to numerous sequencing experiments of IBV (especially vaccine viruses). This protocol demonstrates the feasibility of real-time-based sequencing to provide isolate-level resolution of IBV directly from clinical swab samples, including samples that contain multiple isolates.

The proper diagnosis of IBV as the cause of clinical respiratory disease is contingent on virus typing and differentiating live vaccine viruses from field strains.^11^ Previously, it was reported that IBV genotypes correlate well with the serotypes of IBV^45^; therefore, accurate genotypic identification of IBV will be useful to diagnose vaccine and variant viruses in clinical samples. The rapid pan-IBV and serotype specific RT-qPCR assays^4,23,36^ have been used for serotype identification; however, positive results from RT-qPCR is insufficient to determine the IBV genotype and obtain isolate-level resolution of IBV; thus, sequence analysis of IBV-S1 gene is required. Previously, use of partial S1 gene sequences (450 bp) to type IBV has been described.^29^ However, increasing the length of sequenced S1 gene (~1620 bp) results in more data to be used for genotyping as more of the hypervariable region is covered.^17,27,45^ For example, the AmpSeq protocol was able to differentiate two genetically very similar (99.5% at S1 gene of IBV genome) but serotypically different GIV-L1 isolates, DE072 and GA98, within a single sample. Thus, the detection of highly diverse IBV genotypes, via the S1 subunit, with a single and rapid sequencing protocol is desirable and the ability to detect multiple isolates in a single sample would improve IBV detection in clinical samples.

Currently, pan-IBV RT-qPCR is used to rapidly detect IBV from clinical samples for screening purposes,^4^ but this requires additional serotype-specific RT-qPCRs to genotype positive samples, including samples containing more than one genotype. Additionally, if unidentified IBVs are in the sample, then current detection and characterization may also require egg culture followed by various PCRs.^8^ The AmpSeq method described in this report detected IBV in 13 of 14 IBV-positive and detected all the mixed isolates in four of seven samples containing two or more isolates. While mixed isolates were not 100% identified (0/2 in #1, 2/4 in #2, and ½ in #3) in three samples with the AmpSeq method, detection of those mixed isolates required several non-AmpSeq assays. Although, there is room for improving this new AmpSeq method, it represents a promising, single-step assay that can be used without egg culture. Thus, it is another tool for the detection of IBV, especially in cases when multiple isolates may be present or when genotyping is especially important.

One area that is problematic for a test such as AmpSeq is to detect all the isolates that may be present. While AmpSeq did detect all isolates in 4 samples, in 3 cases not all of the isolates were detected. A potential explanation to partial detection of multiple IBV genotypes could be the relative abundance of IBV genotypes in these clinical samples. It could also be that amplification of these IBV genotypes by serotype-specific RT-qPCR assays is more efficient due to small targeted fragment size (e.g., 120 bp for PCR and ~1600 bp for AmpSeq) and better primer alignment to the target (e.g., degenerate bases are used in the S1 primers used for AmpSeq). One complicating factor of the current AmpSeq protocol is that the IBV target sequence^27^ was not originally designed for high specificity, especially from clinical samples. As such, a high proportion of total reads were non-IBV reads (e.g., often mapping to the chicken genome, data not shown), consistent with the extra bands visible in the original report for these target sequences.^28^ Additionally, it is possible that certain genotypes are better complemented to the S1 primers than others and may outcompete those isolates for amplification in the AmpSeq protocol. Lastly, increasing the total number of reads collected by AmpSeq may improve the ability to detect all genotypes in sample by AmpSeq. It is possible to allow the sequencing to continue longer to obtain more reads per sample. Overall, AmpSeq is a feasible test for IBV characterization, and work is ongoing to improve this new type of assay.

For example, cost and time efficiency of a sequencing protocol can be improved through multiplexing of more samples in a single sequencing run.^3,47^ In this study, samples (Run 1 = 10, Run 2 = 5) were simultaneously multiplexed (i.e., pooled and then sequenced in one run) while maintaining IBV genotyping from data collected. Since the MinION flow cells were not exhausted, and can be washed and re-used, the AmpSeq method also has the potential for sequential multiplexing. This would decrease the need to hold samples for weeks while waiting for the cost-optimal number of samples for simultaneous multiplexing. Alternatively, if detection of all isolates within a given sample is preferred, the sequencing can be run longer to increase sensitivity. The single protocol nature of AmpSeq, the ability to obtain S1 gene sequence results, real-time data analysis, and flexibility of testing design makes MinION-based AmpSeq a viable sequencing protocol for typing of IBV isolates.

The advent of real-time, in-house, third generation sequencing represents a transformative opportunity for diagnostic laboratories, by offering the ability to more fully characterize PCR reactions beyond confirming amplicon size (e.g., routine electrophoresis), Sanger sequencing RT-PCR products, or by confirming a partial sequence through probe hybridization (e.g., probe-based qPCR). However, the interpretation of such large data sets represents a challenge to veterinary diagnosticians. Read-based classification software such as Centrifuge^24^, Kraken^50^, QIIME^5^ and Mothur^37^ have been used to identify and profile microbial species; however, the high error rate^20^ of Nanopore reads translates to poor classification accuracy for many of these tools. Alternatively, *de novo^3,26^* or reference-based^18^ assembly methods have been used in MinION and other deep sequencing platforms. Using a strategy similar to other unbiased laboratory tests (e.g., standard bacterial cultures), an approach was developed to maximize usage of reads (i.e., reads are not discarded based on pre-set length or abundance requirements, similar to how a single colony may be interpreted as a significant result). This approach uses read-based classification against a database containing an equal number of representative sequences per lineage to detect and classify the reads based on their lineage, before conducting read-based classification to the isolate level by using all available IBV sequences in a separate database. Finally, lineage-clustered isolate-level read assignments are interpreted by a veterinary diagnostician to result in reads available for consensus building. The use of a final consensus alignment helps to overcome the individual error rate of MinION sequencing.^3^ Similarly, the absence of predetermined metrics used in de novo assembly allows for the informed decision as to how many consensus sequences to build, a bioinformatics problem when dealing with clinical cases that can contain more than one isolate (e.g., similar to how there is not a predetermined number of significant bacterial colonies). Confirmatory follow-up tests (e.g., RT-PCR) may be needed when dealing with low number of reads in clinical samples; however, future testing of this new technology will allow for creating standards for such confirmatory testing.

Taken together, these results suggest the application of MinION-based AmpSeq, specifically for detecting IBV isolates from clinical samples within a few days, compared to several days to weeks and multiple diagnostic assays to culture and detect multiple IBV genotypes from single sample. Thus, the MinION-based AmpSeq coupled with the data analysis workflow for identification, differentiation, and accurate prediction of IBV genotypes from clinical swab samples can be used as an adjunct to other established rapid diagnostic assays^9,12^ until extensive testing of this protocol is done to improve and validate AmpSeq for IBV diagnostics. Furthermore, AmpSeq-based assays can be, and are being applied to other viral pathogens,^3,37^ demonstrating the power and utility of this method in the age of molecular diagnostics.

## Acknowledgements

We acknowledge Vanessa Gauthiersloan, Samantha Day and Erich Linnemann from the Poultry Diagnostic & Research Center, University of Georgia for the technical help.

## References

1. Abro, SH. Emergence of novel strains of avian infectious bronchitis virus in Sweden. Vet Microbiol 2012;155:237–246.

2. Bochkov, YA. Phylogenetic analysis of partial S1 and N gene sequences of infectious bronchitis virus isolates from Italy revealed genetic diversity and recombination. Virus Genes 2007;35:65–71.

3. Butt, SL. Rapid virulence prediction and identification of Newcastle disease virus genotypes using third-generation sequencing. Virol J 2018;15:179.

4. Callison, SA. Development and evaluation of a real-time Taqman RT-PCR assay for the detection of infectious bronchitis virus from infected chickens. J Virol Methods 2006;138:60–65.

5. Caporaso, JG. QIIME allows analysis of high-throughput community sequencing data. Nat Methods 2010;7:335.

6. Casais, R. Gene 5 of the avian coronavirus infectious bronchitis virus is not essential for replication. J Virol 2005;79:8065–8078.

7. Cavanagh, D. Coronavirus avian infectious bronchitis virus. Vet Res 2007;38:281–297.

8. Cavanagh, D. Infectious bronchitis. Diseases of poultry 2003;11:101–119.

9. Chousalkar, K. LNA probe-based real-time RT-PCR for the detection of infectious bronchitis virus from the oviduct of unvaccinated and vaccinated laying hens. J Virol Methods 2009;155:67–71.

10. De Groot, R. Evidence for a coiled-coil structure in the spike proteins of coronaviruses. J Mol Biol 1987;196:963–966.

11. De Herdt, P. Infectious bronchitis virus infections of chickens in Belgium: an epidemiological survey. Vlaams Diergeneeskundig Tijdschrift 2016;85:285–290.

12. Escutenaire, S. SYBR Green real-time reverse transcription-polymerase chain reaction assay for the generic detection of coronaviruses. Arch Virol 2007;152:41–58.

13. Ferreira, H. Presence of Newcastle disease viruses of sub-genotypes Vc and VIn in backyard chickens and in apparently healthy wild birds from Mexico in 2017. Virus Genes 2019.

14. Hodgson, T. Neither the RNA nor the proteins of open reading frames 3a and 3b of the coronavirus infectious bronchitis virus are essential for replication. J Virol 2006;80:296–305.

15. Jackwood, MW. Molecular evolution and emergence of avian gammacoronaviruses. Infect Genet Evol 2012;12:1305–1311.

16. Jackwood, MW. Data from 11 years of molecular typing infectious bronchitis virus field isolates. Avian Dis 2005;49:614–618.

17. Jackwood, MW. Further development and use of a molecular serotype identification test for infectious bronchitis virus. Avian Dis 1997:105–110.

18. Jain, M. Improved data analysis for the MinION nanopore sequencer. Nature methods 2015;12:351.

19. Jain, M. The Oxford Nanopore MinION: delivery of nanopore sequencing to the genomics community. Genome Biol 2016;17:239.

20. Jain, M. MinION Analysis and Reference Consortium: Phase 2 data release and analysis of R9. 0 chemistry. F1000Research 2017;6.

21. Johnson, MA. A recombinant fowl adenovirus expressing the S1 gene of infectious bronchitis virus protects against challenge with infectious bronchitis virus. Vaccine 2003;21:2730–2736.

22. Kamble, NM. Evolutionary and bioinformatic analysis of the spike glycoprotein gene of H120 vaccine strain protectotype of infectious bronchitis virus from India. Biotechnol Appl Biochem 2016;63:106–112.

23. Keeler, CL, Jr. Serotype identification of avian infectious bronchitis virus by RT-PCR of the peplomer (S-1) gene. Avian Dis 1998;42:275–284.

24. Kim, D. Centrifuge: rapid and sensitive classification of metagenomic sequences. Genome Res 2016.

25. Kingham, B. Identification of avian infectious bronchitis virus by direct automated cycle sequencing of the S-1 gene. Avian Dis 2000:325–335.

26. Koren, S. Canu: scalable and accurate long-read assembly via adaptive k-mer weighting and repeat separation. Genome Res 2017:gr. 215087.215116.

27. Lee, C-W. Redesign of primer and application of the reverse transcriptase-polymerase chain reaction and restriction fragment length polymorphism test to the DE072 strain of infectious bronchitis virus. Avian Dis 2000:650–654.

28. Lee, C-W. Typing of field isolates of infectious bronchitis virus based on the sequence of the hypervariable region in the S1 gene. J Vet Diagn Invest 2003;15:344–348.

29. Lee, C-W. Evidence of genetic diversity generated by recombination among avian coronavirus IBV. Arch Virol 2000;145:2135–2148.

30. McKinley, ET. Avian coronavirus infectious bronchitis attenuated live vaccines undergo selection of subpopulations and mutations following vaccination. Vaccine 2008;26:1274–1284.

31. Naguib, MM. New real time and conventional RT-PCRs for updated molecular diagnosis of infectious bronchitis virus infection (IBV) in chickens in Egypt associated with frequent co-infections with avian influenza and Newcastle disease viruses. J Virol Methods 2017;245:19–27.

32. Okino, CH. Rapid detection and differentiation of avian infectious bronchitis virus: an application of Mass genotype by melting temperature analysis in RT-qPCR using SYBR Green I. J Vet Med Sci 2018;80:725–730.

33. Pohuang, T. Sequence analysis of S1 genes of infectious bronchitis virus isolated in Thailand during 2008–2009: identification of natural recombination in the field isolates. Virus Genes 2011;43:254–260.

34. Quick, J. Multiplex PCR method for MinION and Illumina sequencing of Zika and other virus genomes directly from clinical samples. Nat Protoc 2017; 12:1261.

35. Quick, J. Real-time, portable genome sequencing for Ebola surveillance. Nature 2016;530:228.

36. Roh, H-J. Detection of infectious bronchitis virus with the use of real-time quantitative reverse transcriptase–PCR and correlation with virus detection in embryonated eggs. Avian Dis 2014;58:398–403.

37. Schloss, PD. Introducing mothur: open-source, platform-independent, community-supported software for describing and comparing microbial communities. Appl Environ Microbiol 2009;75:7537–7541.

38. Spaan, W. Coronaviruses: structure and genome expression. J Gen Virol 1988;69:2939–2952.

39. Spatz, SJ. MinION sequencing to genotype US strains of infectious laryngotracheitis virus. Avian Pathol 2019:1–15.

40. Stenzel, T. Differentiation of infectious bronchitis virus vaccine strains Ma5 and 4/91 by TaqMan real-time PCR. Pol J Vet Sci 2017;20:599–601.

41. Sutou, S. Cloning and sequencing of genes encoding structural proteins of avian infectious bronchitis virus. Virology 1988;165:589–595.

42. Tamura, K. MEGA6: molecular evolutionary genetics analysis version 6.0. Mol Biol Evol 2013;30:2725–2729.

43. Tarnagda, Z. Prevalence of infectious bronchitis and Newcastle disease virus among domestic and wild birds in H5N1 outbreaks areas. The Journal of Infection in Developing Countries 2011;5:565–570.

44. Valastro, V. S1 gene-based phylogeny of infectious bronchitis virus: an attempt to harmonize virus classification. Infect Genet Evol 2016;39:349–364.

45. Wang, C-H. Relationship between serotypes and genotypes based on the hypervariable region of the S1 gene of infectious bronchitis virus. Arch Virol 2000;145:291–300.

46. Wang, J. MinION nanopore sequencing of an influenza genome. Front Microbiol 2015;6:766.

47. Wei, S. Rapid multiplex small DNA sequencing on the MinION nanopore sequencing platform. G3: Genes, Genomes, Genetics 2018:g3. 200087.202018.

48. Wickramasinghe, IA. The avian coronavirus spike protein. Virus Res 2014;194:37–48.

49. Williams, AK. Comparative analyses of the nucleocapsid genes of several strains of infectious bronchitis virus and other coronaviruses. Virus Res 1992;25:213–222.

50. Wood, DE. Kraken: ultrafast metagenomic sequence classification using exact alignments. Genome Biol 2014;15:R46.

51. Ye, J. Primer-BLAST: a tool to design target-specific primers for polymerase chain reaction. BMC Bioinformatics 2012;13:134.

